# Anaerobic Fluorescent Reporters Enable *In Vivo* Tracking of VRE Dynamics in *Enterococcus faecium*

**DOI:** 10.1101/2025.02.08.637206

**Authors:** Benjamin R. Treat, Joseph Hobeika, Milisen Tvarkunas, Cliff Guy, Caitlin E. Billiot, Marygrace Duggar, Trinity Fields, Mollie Black, Christina M. Kohler, Elisa B. Margolis

## Abstract

Vancomycin-resistant *Enterococcus faecium* (VREfm) is a leading cause of healthcare-associated bloodstream infection and a dominant colonizer of the antibiotic-perturbed gut. Experimental interrogation of VREfm colonization dynamics has been constrained by three major barriers: widespread resistance to conventional selection markers in contemporary clinical isolates, instability of plasmid-based constructs, and failure of oxygen-dependent fluorescent reporters in the anaerobic intestinal environment. To address these limitations, we develop genetic tools optimized for VREfm that enable stable anaerobic fluorescent labeling and longitudinal tracking of colonization dynamics. We show that bile pigment–binding fluorescent proteins, including eUnaG2 and smURFP, generate robust signal under anaerobic conditions across diverse clinical backgrounds. Systematic analysis of resistance profiles revealed that conventional selection markers are frequently incompatible with modern isolates; accordingly, we identify puromycin as a broadly effective entry-point strategy and demonstrate that a puromycin-selectable plasmid enables reporter expression for otherwise genetically intractable extensively drug-resistant (XDR) strains. To support long-term studies without continuous selection, we establish an enhanced *pheS*\*\* counterselection system for chromosomal integration at neutral loci and engineer a tandem 2×smURFP construct to optimize single-copy fluorescence. Using these tools, we directly track competition between fluorescently labeled VREfm strains in a murine model via flow cytometric analysis of stool samples, enabling longitudinal strain-level quantification without antibiotic selection or selective plating. This work provides a unified framework for genetic manipulation and anaerobic strain tracking in XDR *E. faecium*, establishing a practical platform for dissecting ecological processes that govern persistence and competition in the gut.

**Importance:** Vancomycin-resistant *Enterococcus faecium* (VREfm) is a dangerous and difficult-to-treat hospital pathogen. How VREfm establishes itself in the gastrointestinal tract and competes with other strains remains poorly understood, yet this knowledge is essential for preventing infection. Studying these processes has been impeded by widespread antibiotic resistance and by the failure of conventional fluorescent proteins in the oxygen-free gut environment. We describe tools that overcome these barriers and provide a practical foundation for mechanistic studies of VREfm gut colonization and persistence.

## Introduction

Vancomycin-resistant *Enterococcus faecium* (VREfm) is a major nosocomial pathogen and a leading cause of healthcare-associated bloodstream infection in immunocompromised patients(1–3). The burden is especially high in patients receiving intensive chemotherapy or hematopoietic stem cell transplantation, where microbiota disruption often precedes intestinal domination and invasive disease(2–4). In these settings, gut colonization is both a marker and driver of morbidity, mortality, and hospital transmission(2, 5, 6).

The success of VREfm as a hospital-adapted pathogen reflects a combination of intrinsic resilience and extraordinary genomic plasticity. Modern clinical isolates typically harbor resistance determinants to vancomycin, aminoglycosides, macrolides, fluoroquinolones, and frequently additional classes such as tetracyclines and oxazolidinones(5–9). Beyond antibiotic resistance, VREfm exhibits enhanced tolerance to environmental stress, persistence on abiotic surfaces, and an ability to rapidly adapt to selective pressures imposed by antibiotics and host immunity(9, 10). These features have made VREfm a model organism for studying pathogen adaptation in the hospital ecosystem while simultaneously rendering it experimentally intractable.

A critical reservoir for VREfm is the gastrointestinal tract, where it can persist at high abundance for prolonged periods, transmit between hosts, and re-emerge following antibiotic perturbation. Numerous clinical and experimental studies have demonstrated that intestinal domination by VREfm strongly predicts bloodstream infection and facilitates nosocomial spread(2, 5, 6). As a result, there is substantial interest in understanding the ecological and molecular mechanisms that govern VREfm colonization, persistence, and competition within the gut microbiota(11). Such knowledge is essential for developing strategies aimed at preventing colonization, promoting decolonization, or restoring colonization resistance.

Despite this need, experimental interrogation of VREfm population dynamics *in vivo* has lagged behind that of other enteric pathogens. One major limitation is the lack of robust genetic tools that function reliably in contemporary VREfm clinical isolates. While *Enterococcus* has historically been amenable to genetic manipulation in laboratory-adapted strains, modern clinical isolates frequently exhibit heterogeneous but widespread resistance to antibiotics commonly used as selectable markers in Gram-positive bacteria(9, 11). In many cases, no single conventional marker provides reliable coverage across isolates, and iterative genetic manipulation becomes impractical or impossible. This challenge is compounded in extensively drug-resistant (XDR) strains, where the remaining antibiotic susceptibilities are often limited to compounds unsuitable for routine laboratory selection.

Plasmid instability is a second barrier. Many clinical isolates fail to maintain engineered plasmids without continuous selection, which is incompatible with coculture and long-term *in vivo* studies. Chromosomal integration can solve this problem, but existing counterselection systems often perform poorly in *E. faecium*, particularly in the rich media needed for clinical isolates(12).

A third and equally significant barrier is the reliance of most fluorescent and bioluminescent reporters on molecular oxygen. Green fluorescent protein and its derivatives require oxygen for chromophore maturation, while luciferase-based systems depend on oxygen for light production(13, 14). As a consequence, these reporters perform poorly or fail entirely in the anaerobic environment of the intestinal lumen, where VREfm resides(15). This limitation has forced many studies to rely on indirect readouts of colonization, such as selective plating or sequencing-based relative abundance estimates, which lack temporal resolution, are labor-intensive, and provide limited insight into strain-level population dynamics.

Recent efforts have begun to explore oxygen-independent fluorescent proteins and alternative selection strategies in anaerobic bacteria(16–20), but their application to VREfm—particularly across diverse XDR clinical isolates and *in vivo* settings—remains limited. Moreover, existing approaches have rarely been integrated into a unified toolkit capable of supporting both genetic manipulation and ecological measurements such as strain competition.

To address these challenges, we developed a set of genetic tools optimized specifically for XDR *E. faecium*. This toolkit integrates three core capabilities: (i) a universal entry-point selection strategy that enables initial genetic access to otherwise intractable clinical isolates, (ii) engineered anaerobic fluorescent reporters that function in anaerobic environments such as the gut, and (iii) robust chromosomal integration using improved counterselection at neutral genomic loci to support long-term *in vivo* studies. Importantly, these tools are designed not only for strain labeling but also to enable direct measurement of ecological processes, including *in vivo* strain competition, without reliance on antibiotic markers. In this study, we describe the development and validation of these tools and demonstrate their utility in a murine model of VREfm intestinal colonization.

## Results

### Engineering Oxygen-Independent Fluorescent Labeling in Clinical VREfm Isolates

Because oxygen-dependent reporters perform poorly in the intestinal lumen(13–15), we tested whether oxygen-independent systems could support strain-resolved fluorescence in VREfm under anaerobic conditions.

To enable fluorescence in *Enterococcus faecium*, we engineered a modular shuttle plasmid platform optimized for oxygen-independent labeling in clinical VREfm isolates (Figure 1A). This E. coli shuttle backbone includes an Enterococcus replication origin (pAMβ1), the pCF10 oriT which allows efficient *E. faecalis* CK111 conjugative transfer, and chloramphenicol and spectinomycin resistance markers to allow for broad isolate compatibility. Reporter genes were placed under the constitutive *S. agalactiae cfb* promoter, a strong promoter derived from the CAMP factor locus and previously described for high-level expression in Gram-positive bacteria(21).

**Figure 1.**
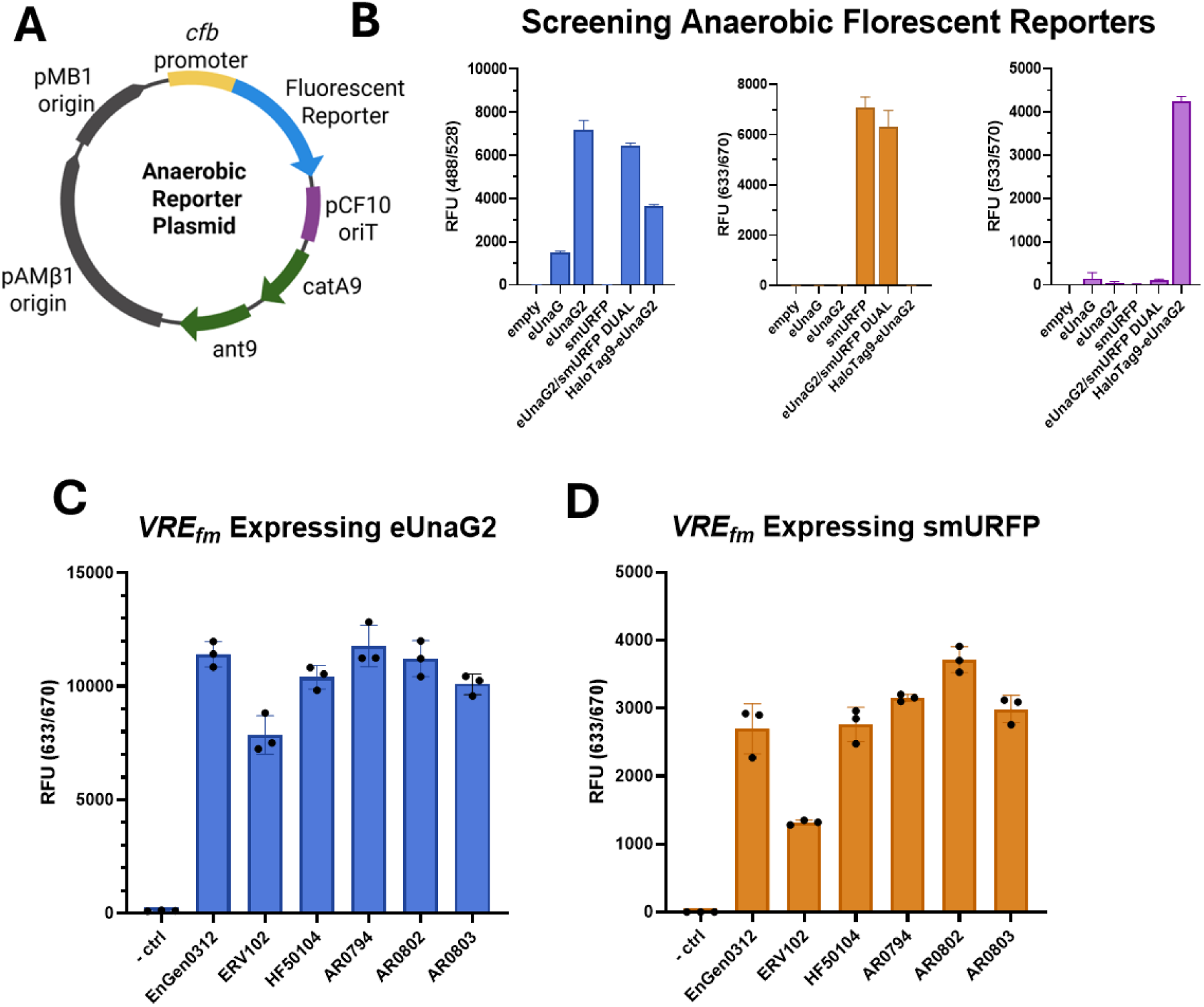
Anaerobic Fluorescent Reporters designed for VRE *E. faecium.* (**A**) Anaerobic Reporter Plasmid includes an E. coli origin (pMB1), an Enterococcus origin (pAMβ1), the S. *agalactiae cfb* promoter, anaerobic fluorescent reporter genes (eUnaG, eUnaG2, smURFP, eUnaG2/smURFP dual expression, or a HaloTag9-eUnaG2 fusion gene), the pCF10 origin of transfer (oriT), and antibiotic resistance markers (catA9/chloramphenicol resistance, ant9/spectinomycin resistance). (**B**) Relative fluorescence of reporter plasmids in conjugal donor strain *E. faecalis* CK111. Two independent transformants are represented for each group. (**C**) and (**D**) Relative fluorescence of VREfm strains transformed with eUnaG2 or smURFP reporter plasmids. Fluorescence measures performed after overnight growth in BHI supplemented with 5nM biliverdin and 25nM bilirubin. Cultures were centrifuged at 6,000g and resuspended in 1XPBS prior to fluorescence readings on a BioTek Synergy H1 microplate reader. For fluorescence detection, 2-4 independent transformants were analyzed. Data plotted as mean with SD in Graphpad Prism 10.4. Plasmid schematic created with BioRender.com.

We focused on bile pigment–binding fluorescent proteins, which generate signal independently of oxygen through ligand binding rather than oxidative chromophore maturation. Specifically, we evaluated eUnaG2(18), an engineered derivative of UnaG(16, 17) optimized for brightness and stability, and smURFP(19), a monomeric biliverdin-binding far-red fluorescent protein previously shown to function in low-oxygen and deep-tissue environments. In parallel, we evaluated HaloTag9(20), a self-labeling enzyme that requires covalent attachment of a synthetic fluorophore ligand (e.g., HaloTag® TMR Ligand; many other HaloTag-compatible ligands are available) but would provide spectral flexibility.

Several configurations were constructed for testing in VREfm: eUnaG, eUnaG2, smURFP, and a HaloTag9–eUnaG2 fusion, along with a dual smURFP/eUnaG2 construct. We first transformed the *E. faecalis* CK111 conjugal donor strain, where eUnaG2 and smURFP produced strong, reproducible fluorescence, whereas eUnaG was dimmer (Figure 1B). The HaloTag fusion generated bright signal following ligand addition but required exogenous labeling and careful washing steps.

Reporter plasmids were next introduced into multiple VREfm strains spanning distinct clinical lineages. Under strictly anaerobic overnight growth in BHI supplemented with nanomolar bilirubin or biliverdin, both eUnaG2- and smURFP-expressing strains produced strong and readily detectable fluorescence by microplate reader (Figure 1C–D). Signal separation between labeled and unlabeled populations was robust across independent transformants and strain backgrounds. Although fluorescence magnitude varied modestly between isolates—likely reflecting differences in plasmid copy number, promoter activity, or intracellular ligand availability—strain-specific differences did not abrogate reporter function.

Unlike HaloTag9, eUnaG2 and smURFP require no enzymatic labeling or wash steps; fluorescence arises from spontaneous bile-pigment binding and may therefore occur in the intestine without exogenous chromophore addition. These reporters provided reliable anaerobic labeling across diverse VREfm isolates. During this work, however, we found that conventional selectable markers were often ineffective in XDR backgrounds, prompting the selection analysis below.

### Puromycin establishes a universally effective selection marker for XDR VREfm

To define how antibiotic resistance limits genetic manipulation of contemporary *E. faecium*, we surveyed resistance profiles in vancomycin-resistant *Enterococcus* isolates from the CDC AR Isolate Bank(22). The panel included nine *E. faecium* isolates plus representative *E. faecalis*, *E. casseliflavus*, and *E. lactis* strains, and covered antibiotics commonly used as Gram-positive selection markers.

Resistance to conventional selection agents was widespread but heterogeneous (Figure 2A and Supplemental Figure 1B). High-level aminoglycoside resistance was common(8, 9), macrolide and tetracycline resistance were frequent(7, 9), and chloramphenicol resistance was observed in multiple strains(9).

**Figure 2.**
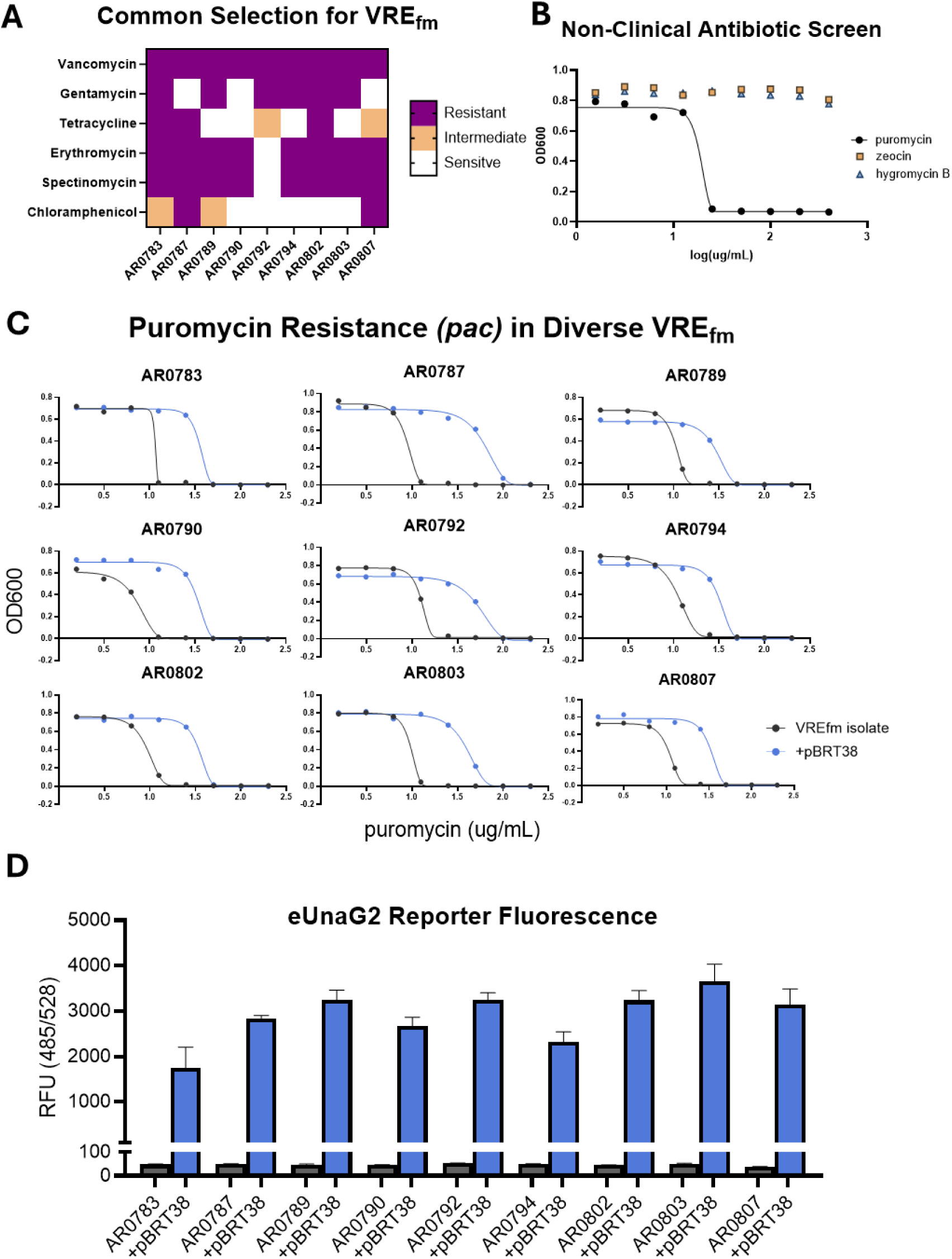
Universal XDR VREfm Labeling via Puromycin-Resistance Marker. (**A**) Heatmap of resistances to many common antibiotics for selection in Enterococci. All Isolates obtained from the CDC VRE AR Isolate Bank. Purple indicates high-level resistance, orange intermediate resistance, and white indicates sensitivity. (**B)** Minimum Inhibitory Concentration (MIC) assays of puromycin, zeocin, and hygromycin B were performed on VREfm strain TX0082. (**C**) MIC assays for puromycin of 9 VREfm strains from CDC AR Isolate Bank, wild-type are in blue, and transformants expressing *pac* from pBRT38 are in black. (**D**) Relative fluorescence of VREfm strains transformed with puromycin-resistant eUnaG2 reporter plasmids. Fluorescence measures performed after overnight growth in BHI supplemented with 25nM bilirubin.

While individual isolates retained susceptibility to one or more agents, no single conventional marker provided reliable coverage across the full isolate panel. In practice, this heterogeneity necessitates strain-specific selection strategies and complicates comparative or iterative genetic engineering, particularly in extensively drug-resistant (XDR) backgrounds where remaining antibiotic susceptibilities are limited.

We therefore tested three nonclinical laboratory antibiotics: puromycin(23–25), hygromycin B(26), and zeocin(27). By broth microdilution, puromycin consistently inhibited all isolates (Figure 2B), with wild-type MICs clustering in a narrow 8 to 16 µg/mL range (Supplemental Table 1).

To determine whether puromycin resistance could be genetically conferred, we constructed plasmid pBRT38 expressing a codon-optimized puromycin N-acetyltransferase (*pac*) gene alongside the eUnaG2 reporter (Supplemental Figure 1A). Expression of *pac* reproducibly increased puromycin MICs approximately 3- to 8-fold across all nine CDC *E. faecium* isolates tested (Figure 2C; Supplemental Table 1). Transformants maintained strong eUnaG2 fluorescence under puromycin selection (Figure 2D), demonstrating compatibility between selection and anaerobic reporter expression. In contrast, zeocin and hygromycin B exhibited variable activity and failed to provide consistent inhibition across isolates (Figure 2B), limiting their utility as broadly applicable selection markers in XDR Enterococcus.

Together, these data establish puromycin as a universally effective entry-point selection strategy for genetically intractable VREfm isolates. However, the relatively narrow selection window observed across strains suggested that puromycin would be best suited for multicopy plasmid-based applications rather than single-copy chromosomal insertions, motivating the chromosomal integration strategy described below.

### In Vivo Detection of VREfm and Limitations of Plasmid-Based Labeling

We next asked whether plasmid-encoded reporters could be detected in the murine intestine (Figure 3). Mice pretreated with vancomycin were inoculated with VREfm expressing plasmid-borne smURFP, and stool collected 24 h later contained a distinct fluorescent bacterial population detectable by direct flow cytometry without enrichment or selective plating (Figure 3A). Whole-stool imaging and intestinal confocal microscopy confirmed *in vivo* reporter signal (Figure 3B and C).

**Figure 3.**
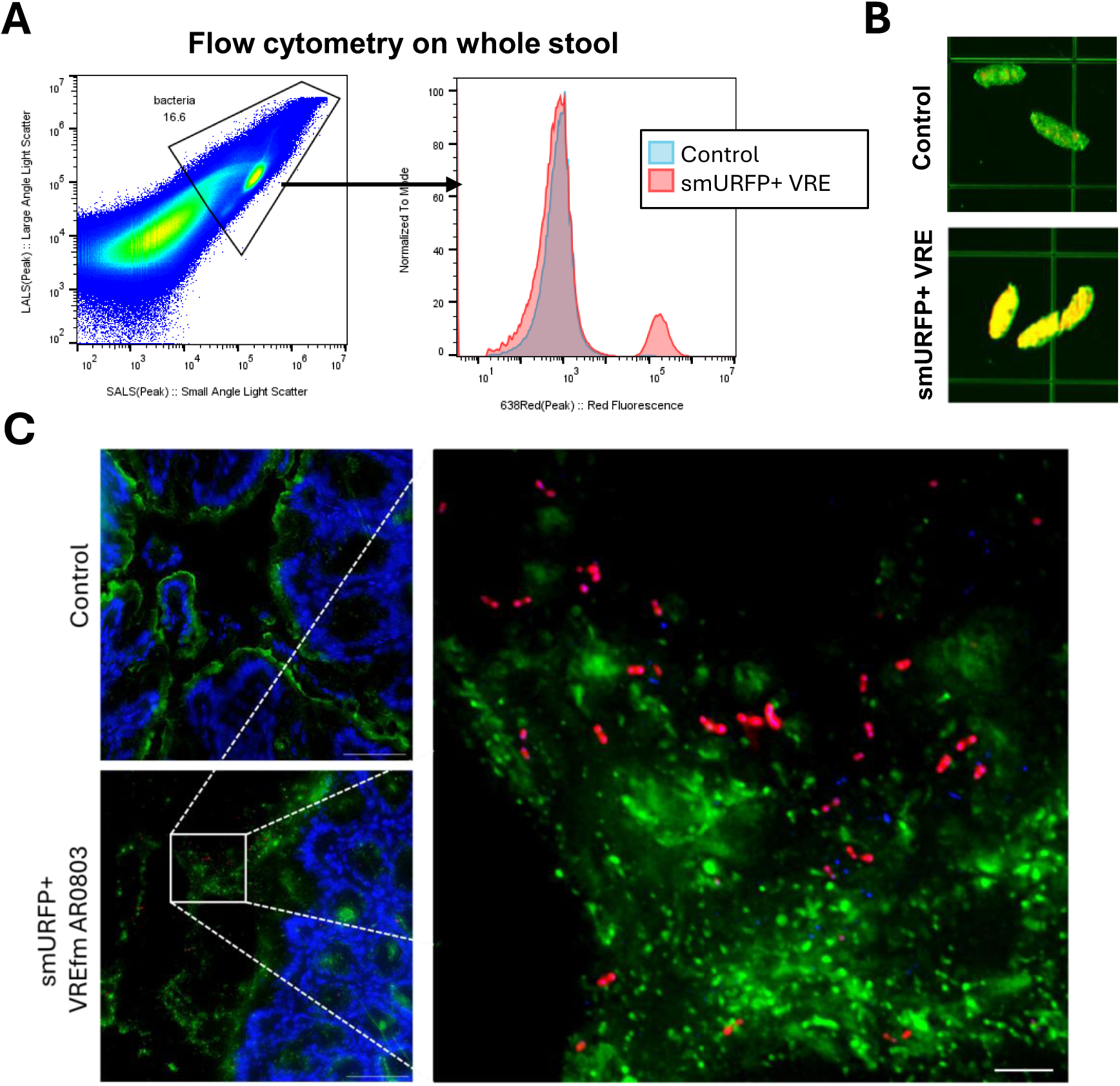
Tracking Anaerobic VREfm Fluorescence *in vivo*. 6-8 week old female C57BL/6 mice (Taconic Biosciences) were placed on Vancomycin (0.5mg/mL in water) for 48 hours prior to inoculation with 1×10^8^ CFU *E. faecium* AR0803 containing pBRT31 (smURFP reporter). (**A**) Fresh stool from mice was collected 24 hours post inoculation (hpi), resuspended in PBS at 10% w/v, filtered through a 40 µm strainer, and analyzed on an ApogeeFlow MicroPLUS. Events were gated broadly on the total bacterial population using SALS and LALS parameters before smURFP fluorescence gating. (**B**) Whole stool fluorescence at 24 hpi (BioRad ChemiDoc MP, green indicates 488nm with 532/28 filter, red indicates 647nm with 700/50 filter) (**C**) Confocal microscopy of 10μm unfixed intestinal sections stained with smURFP in red, wheat germ agglutinin (WGA) AF555 in green and DAPI in blue.

Although plasmid-based reporters enabled rapid *in vivo* validation, we observed substantial variability in plasmid retention across clinical VREfm isolates in the absence of continuous antibiotic selection. Serial passaging of smURFP-expressing strains without selection resulted in marked plasmid loss in multiple CDC AR isolates (Supplemental Figure 2), indicating that multicopy constructs are not stably maintained across backgrounds. Because continuous antibiotic selection is incompatible with many *in vivo* and competition experiments, these findings highlighted the need for a chromosomal integration strategy to achieve stable, selection-independent reporter expression. We therefore next developed a genomic insertion framework compatible with XDR VREfm isolates.

### Genomic Integration and Counterselection Enable Stable Anaerobic Reporter Labeling

For long-term *in vivo* studies, we adapted phenylalanine sensitivity-based counterselection using an *E. faecalis pheS* allele previously used in other Gram-positive systems(12, 28). Because standard *pheS*\* performed poorly in clinical VREfm, we incorporated mutations guided by *E. coli* optimization(29) to generate *pheS*\*\* (T258A/A312G), which improved double-crossover recovery in BHI and also worked in additional *Enterococcus* species.

To enable stable chromosomal labeling without disrupting native gene function, we next identified neutral genomic loci suitable for reporter integration. Based on our survey of clinical genomes spanning Clade A *E. faecium* and Clade B isolates (now reclassified as *E. lactis*), we prioritized conserved intergenic regions lacking annotated coding sequences (preferentially between convergent genes) and predicted to tolerate insertions. Two candidate loci—between ORFs 675/680 and 12650/12655—met these criteria and were selected for testing (Figure 4A). The 675/680 intergenic region lies between *gshF* and *yueI* and showed high transcriptional activity(30), whereas the 12650/12655 locus lies between an L,D-transpeptidase and *yebC* and showed moderate transcriptional activity(30).

**Figure 4.**
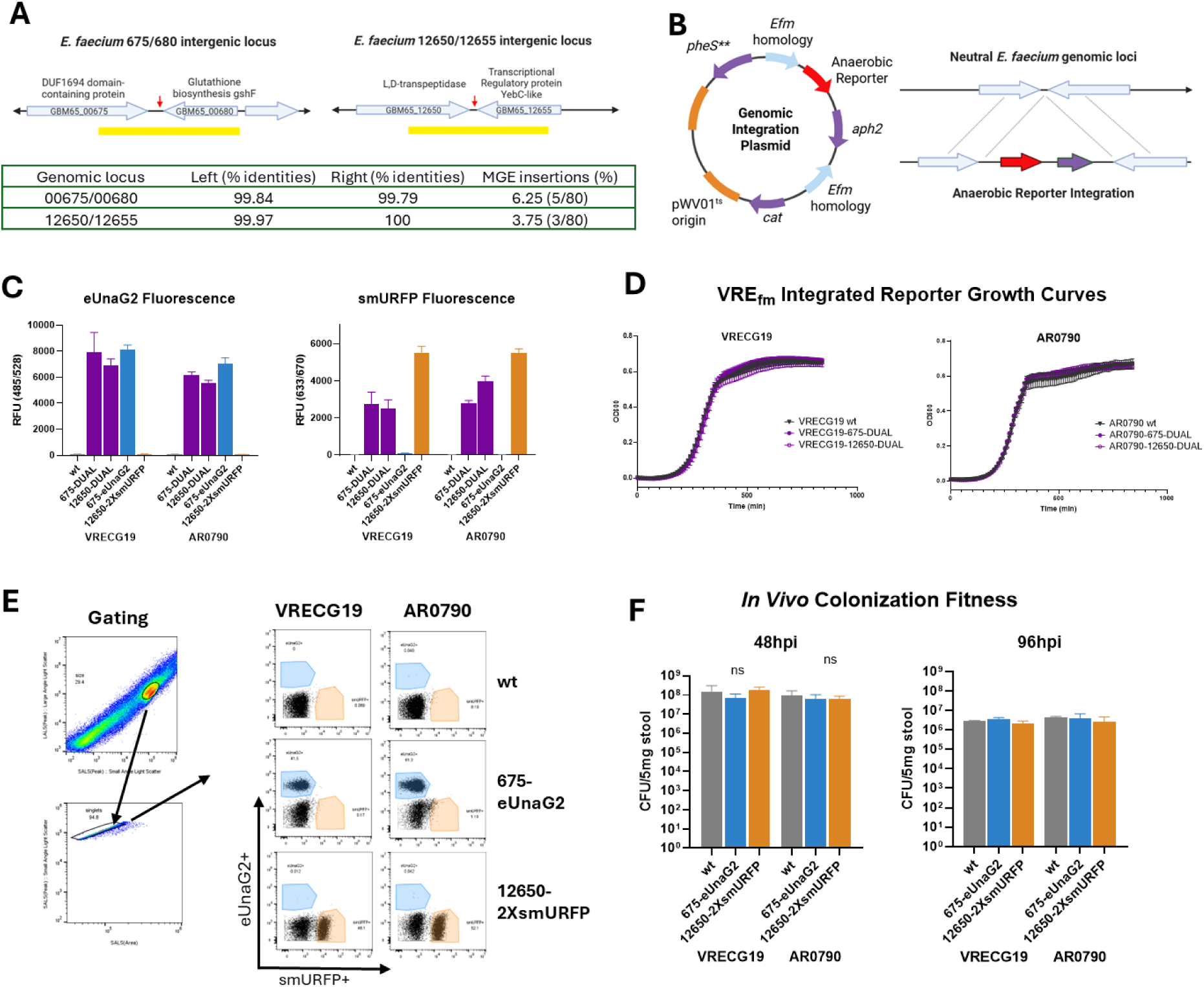
Neutral Genomic Integration Sites in E. faecium. (**A**) Schematic of the 675/680 and 12650/12655 chromosomal insertion loci in E. faecium VRECG19-10-S (GenBank: WEFX01000001.1). The 675/680 locus lies between gshF and yueI, and the 12650/12655 locus lies between an L,D-transpeptidase and yebC. Yellow bars indicate homology regions used for genomic insertion plasmids, and red arrows indicate naturally occurring MGE insertions observed across surveyed clinical genomes. (**B**) Plasmid maps and recombination schematics for genomic insertion plasmids pBRT48 and pBRT63. Plasmids contain: (1) homology arms targeting the 675/680 locus (pBRT48) or 12650/12655 locus (pBRT63), (2) anaerobic fluorescent reporter genes, (3) gentamicin resistance, (4) the pWV01 temperature-sensitive origin, (5) chloramphenicol resistance (cat), and (6) pheS** counterselection for plasmid loss. (**C**) eUnaG2 and smURFP fluorescence reporter activity of two VREfm strains (VRECG19 and AR0790) with single-color or dual genomic insertions in the indicated loci. Overnight growth in BHI supplemented with 5nM biliverdin and 25nM bilirubin. Cultures were centrifuged at 6,000g and pellets resuspended in 1xPBS prior to fluorescence readings on a BioTek Synergy H1 microplate reader. (**D**) Overnight growth curves of wild-type, 675-DUAL, or 12650-DUAL genomic insertions of two VREfm strains (VRECG19 and AR0790). Growth curves were performed in a 96-well plate in triplicate on a BioTek Synergy H1 microplate reader overnight at 37°C in an anaerobic chamber (Coy Laboratory Products). Data plotted as mean with SD in Graphpad Prism 10.4. (**E**) Female C57BL/6 mice (6–8 weeks; n=5/group, Taconic Biosciences) received vancomycin (0.5 mg/mL in drinking water) for 2 days before inoculation with 1×104 CFU E. faecium reporter strains. Stool samples were collected at 48 hpi and resuspended in 1xPBS (10% w/v) containing biliverdin (2.5 nM) and bilirubin (12.5 nM), incubated on ice for 30 min, centrifuged through a 40 µm mesh filter, and resuspended in 1xPBS. Flow cytometry was performed on an ApogeeFlow Micro-PLUS. (**F**) Stool collected at indicated days post-inoculation (dpi) was resuspended in 1xPBS (10% w/v), serially diluted, and plated on BHI agar supplemented with vancomycin (30 µg/mL), meropenem (20 µg/mL).

Plasmid maps and recombination schematics for integration vectors targeting each locus are shown in Figure 4B. To support both benchmarking and downstream competition studies, we generated chromosomal insertions carrying either a dual eUnaG2/smURFP cassette or single-color reporters (eUnaG2 or tandem 2×smURFP). To assess strain generalizability, insertions were introduced into two divergent VREfm backgrounds—VRECG19 (Clade A1, VanA) and AR0790 (Clade A2, VanB). Chromosomal reporter activity was confirmed across both strains, and inclusion of single-color derivatives demonstrated clear signal separation for both eUnaG2 and 2×smURFP (Figure 4C).

Chromosomal integration did not measurably impair growth: wild-type and dual-reporter strains showed indistinguishable anaerobic growth curves in both backgrounds (Figure 4D). *In vivo* detection of VRECG19-DUAL confirmed stable chromosomal expression but also showed that single-copy smURFP was too dim for robust flow-cytometric separation (Supplemental Figure 4). We therefore engineered a tandem 2×smURFP construct using independently codon-optimized ORFs to improve brightness while minimizing recombination. Single-color 675-eUnaG2 and 12650-2×smURFP strains were readily detected *in vivo* and showed no colonization defect relative to parental strains at 48 and 96 h (Figure 4E and F).

### *In vivo* tracking of VREfm strain competition using anaerobic fluorescent reporters

To maximize signal intensity and enable single-copy chromosomal labeling across strain backgrounds, we modified our genomic insertion plasmids to express either eUnaG2 alone or tandem 2×smURFP. These configurations increased reporter brightness and improved utility for competition assays while maintaining compatibility with neutral chromosomal integration.

We next asked whether chromosomally encoded anaerobic reporters could resolve competitive dynamics between VREfm strains both in vitro and in vivo (Figure 5). *In vitro* benchmarking assays were first performed by mixing equal amounts of VREfm strains carrying genomic insertions of either eUnaG2 or 2×smURFP (10⁴ CFU per strain) in BHI supplemented with biliverdin and bilirubin. After incubation, fluorescence values were normalized to OD₆₀₀, scaled relative to single-color controls, and used to calculate a relative competition index (Figure 5A–B). Across multiple isolate pairings, this approach reproducibly detected strain-dependent differences in competitive performance, confirming that reporter output accurately reflects relative abundance in mixed populations.

**Figure 5.**
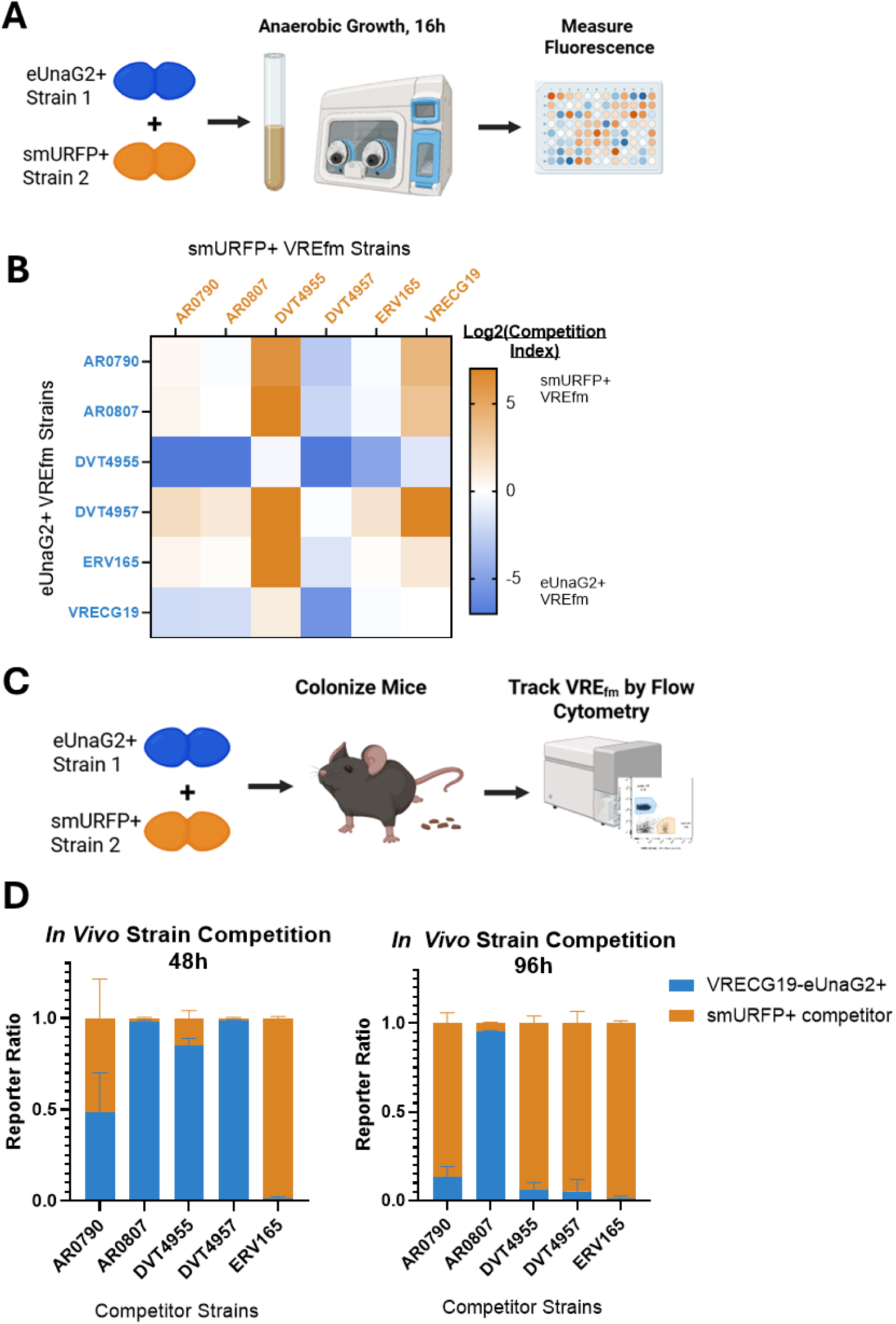
Anaerobic Fluorescent Reporters Allow Tracking of VRE_fm_ Strain Competition. (**A**) Experimental setup for anaerobic VREfm strain competition assays. VREfm strains containing genomic insertions of either eUnaG2 or 2×smURFP were mixed at equal amounts (10⁴ CFU per strain) in BHI supplemented with biliverdin (5 nM) and bilirubin (25 nM) in a total volume of 200 µL in 96-well plates. Plates were centrifuged to pellet cells, resuspended in 1× PBS, and fluorescence measured after 16 h. (**B**) Six isolates expressing either eUnaG2 or 2×smURFP were competed as described above. Fluorescence values were normalized to OD₆₀₀ and scaled to single-color controls to estimate relative abundance. Competitive outcomes are shown as a relative competition index, calculated as log₂(smURFP/eUnaG2); positive values indicate smURFP strain dominance and negative values indicate eUnaG2 strain dominance. Color scale is capped at ±7 for visualization. (**C**) Experimental setup for *in vivo* VREfm strain competition. 6-8 week old female C57BL/6 mice (Taconic Biosciences) were placed on Vancomycin (0.5mg/mL in water) for 48 hours prior to inoculation. Mice were gavaged with VREfm strains containing genomic insertions of either eUnaG2 or 2×smURFP were mixed at equal amounts (10⁴ CFU per strain in 100 µL total 1xPBS). (**D**) Stool samples were processed and analyzed by flow cytometry as previously to quantify the relative abundance of eUnaG2+ and smURFP+ VREfm.

We next applied the system in mice. After vancomycin pretreatment and withdrawal, animals were gavaged with equal mixtures of chromosomally labeled strains, and stool was analyzed longitudinally by direct flow cytometry without enrichment or selective plating (Figure 5C and Supplemental Figure 5). Distinct eUnaG2+ and smURFP+ populations were readily resolved, revealing dynamic, strain-dependent shifts over time that were not always predicted by overnight *in vitro* assays.

This approach enabled repeated within-host sampling without antibiotic selection or invasive procedures. Fluorescent populations remained detectable after freezing, and minority populations could be resolved even when strains differed substantially in abundance. Together, these data show that anaerobic fluorescent reporters support direct, longitudinal measurement of VREfm competition *in vivo*.

## Discussion

Experimental study of intestinal *Enterococcus faecium* ecology has been limited by the same features that make VREfm clinically successful: extensive drug resistance, plasmid instability, and the failure of conventional reporters in anaerobic environments. Here we developed a modular toolkit that addresses these barriers and enables direct strain-level tracking *in vivo*.

A major obstacle is that standard Gram-positive selection markers are poorly matched to the resistance landscape of contemporary VREfm. Our strain panel showed widespread but uneven resistance to aminoglycosides, macrolides, tetracyclines, and chloramphenicol(7, 8, 10, 31), emphasizing that genetic intractability in hospital-adapted lineages is partly a biological consequence of antibiotic adaptation, not just a technical problem. Puromycin provided the broadest entry-point solution. Susceptibility was conserved across tested enterococci, and multicopy *pac* expression supported reliable plasmid selection, consistent with the conserved ribosomal target of puromycin and its lack of clinical use(23–25). That said, the narrow selection window makes puromycin better suited to multicopy plasmids than to the chromosomal workflow used here. Extending broad-compatibility selection to single-copy integration remains an important next step and will likely require stronger expression cassettes or more favorable insertion contexts.

Beyond selection, a second major barrier to studying VREfm ecology has been the failure of conventional reporters under anaerobic conditions or within the intestinal environment. Most fluorescent proteins and bioluminescent systems require molecular oxygen for chromophore maturation or light production, limiting their use for organisms that reside primarily in the gut lumen(13–15). By systematically evaluating oxygen-independent systems, we show that bile pigment–binding fluorescent proteins such as eUnaG2 and smURFP provide robust fluorescence across diverse XDR Enterococcus backgrounds under anaerobic conditions. HaloTag9 offered strong signal and spectral flexibility when labeled appropriately, but its reliance on expensive exogenous ligands and additional processing steps likely limits scalability for large cohorts or longitudinal *in vivo* studies. Notably, reporter performance was largely strain-independent, indicating broad portability across contemporary clinical isolates. However, effective fluorescence in complex environments such as stool may depend on microbiome composition, as other microbial community members can metabolize or degrade biliverdin and bilirubin, potentially limiting chromophore availability for reporter activation, and necessitating exogenous addition of ligands before assays.

During tool optimization, we also observed that promoter performance differed between plasmid and chromosomal contexts. Although the *S. agalactiae cfb* promoter yielded strong fluorescence from multicopy plasmids, single-copy chromosomal integration produced insufficient signal for reliable population discrimination by flow cytometry. Rational engineering of a tandem 2×smURFP construct substantially improved brightness while minimizing recombination risk through the use of independently codon-optimized ORFs. This approach provides a broadly applicable strategy for enhancing reporter signal in anaerobic systems and may be useful in other organisms where chromosomal reporter output is limiting.

In parallel with reporter optimization, we designed all reporter and genomic integration plasmids to include an oriT to facilitate conjugative delivery using the readily transformable donor strain *E. faecalis* CK111. This feature was particularly useful for genetically recalcitrant clinical isolates that were difficult to access by direct transformation. In practice, CK111-mediated conjugations were substantially improved by supplementation with the cCF10 peptide pheromone, which consistently increased transfer efficiency across diverse VREfm backgrounds.

Stable chromosomal integration was essential because plasmid retention varied widely without selection. The improved *E. faecalis pheS*\*\* allele markedly increased double-crossover recovery in VREfm grown in rich media, lowering a major technical barrier to chromosomal engineering and providing a framework that should also be useful for future applications such as scarless allelic exchange or larger genetic screens.

Using this framework, we identified conserved intergenic regions that support stable reporter integration without detectable fitness costs across divergent VREfm lineages. Reporter insertions at these sites produced consistent expression across strain backgrounds and supported long-term in vivo tracking without selection. Together with the improved counterselection system, these results establish a practical foundation for stable chromosomal engineering in contemporary VREfm. In addition, this framework is readily adaptable for scarless allelic exchange, as straightforward cassette redesign enables efficient double-crossover insertion of reporters without residual antibiotic resistance sequences.

The key biological advance is that these tools allow direct longitudinal measurement of strain dynamics in the gut. Chromosomally encoded anaerobic reporters enabled repeated, quantitative tracking of competition in individual mice without selective plating or antibiotic selection. This improves temporal resolution, reduces perturbation, and permits detection of minority VREfm populations within communities that may disproportionately contribute to persistence or relapse(5, 6, 11). The discordance we observed between some *in vitro* and *in vivo* outcomes further underscores the importance of measuring competition directly in the intestinal environment. More broadly, this platform should enable experiments testing how antibiotics, microbiota composition, diet, or host immunity shape VREfm colonization and persistence. Because reporter identity is decoupled from antibiotic resistance, strains can be paired flexibly with less selective confounding.

In summary, this work provides a set of genetic tools that address major obstacles in VREfm research: selection incompatibility, anaerobic reporter limitations, counterselection, and plasmid instability. By combining a broadly compatible plasmid entry-point selection strategy with oxygen-independent fluorescence and a scalable chromosomal integration framework, these tools enable direct experimental interrogation of VREfm population dynamics *in vivo*. Given the central role of intestinal colonization in VREfm pathogenesis and transmission, this work should facilitate mechanistic studies aimed at understanding—and ultimately disrupting—the ecological processes that sustain this pathogen in vulnerable patient populations.

## Materials and Methods

### Bacterial strains

Bacterial strains used in this study included VRE isolates from the CDC Antimicrobial Resistance (AR) Isolate Bank(22) and additional VREfm isolates obtained from BEI (EnGen0312, ERV102, HF50104, TX0082) or previously described St. Jude isolates (VRECG13-10-S, VRECG19-10-S)(32). The *E. faecalis* conjugal donor CK111 was described previously(12). All strains were grown in brain heart infusion (BHI; Difco, BD Biosciences). Selection in enterococci used chloramphenicol 30 µg/mL, spectinomycin 300 to 500 µg/mL, puromycin 20 to 30 µg/mL, and gentamicin 300 µg/mL. *E. coli* Stellar (Takara Bio USA) or NEB 5α (New England Biolabs) cells were used for plasmid propagation with chloramphenicol 50 µg/mL, spectinomycin 100 µg/mL, puromycin 100 µg/mL, or gentamicin 25 µg/mL.

### Plasmid Construction

All plasmids were constructed by In-Fusion cloning (Takara) and maintained in Stellar *E. coli*. Reporter plasmids were based on previously described Gram-positive and *E. faecalis* fluorescent reporter backbones(21, 33) and were modified to include *catA9* and pCF10 oriT for improved compatibility with difficult VREfm strains. Anaerobic reporter genes included UnaG (Addgene 163125), eUnaG2 (Addgene 162457), smURFP (Addgene 80341), and HaloTag9 (Addgene 169324), each placed under control of the *S. agalactiae cfb* promoter. A primer-encoded V2L substitution converted UnaG to eUnaG(17). For dual smURFP/eUnaG2 constructs, a ribosome-binding site was inserted between the two ORFs to enable coexpression from the *cfb* promoter.

For puromycin selection validation, we constructed pBRT38, an *Enterococcus*-*E. coli* shuttle vector expressing codon-optimized pac under control of the catA9 promoter (**Supplemental Figure 1B**). The puromycin resistance *pac* gene was codon optimized for *E. faecium* and obtained from GenScript Biotech.

Genomic insertion plasmids were based on the temperature-sensitive pGPA1 delivery plasmid(34) (Addgene 115476), with transposase and transposon sequences replaced by VREfm homology arms and dual smURFP/eUnaG2 reporter cassettes. The pWV01ts origin was repaired by removing a premature stop codon in copG, and *pheS*\*\* was added to enable efficient counterselection on BHI containing 5 to 10 mM PCPA.

Plasmids generated for this work are listed in **Table 2** and will be made available on Addgene prior to full publication of research.

**Table 1.** Puromycin Selection is Universally Compatible with CDC VRE AR Isolates.

| Isolate | Species | Puromycin MIC (µg/mL) |  |
| --- | --- | --- | --- |
|  |  | wt | +pac |
| AR0783 | *E. faecium* | 11.5 | 46.9 |
| AR0787 | *E. faecium* | 11.5 | 96.2 |
| AR0789 | *E. faecium* | 12.4 | 43.0 |
| AR0790 | *E. faecium* | 12.0 | 47.3 |
| AR0792 | *E. faecium* | 16.0 | 77.7 |
| AR0794 | *E. faecium* | 14.2 | 47.2 |
| AR0802 | *E. faecium* | 14.1 | 45.1 |
| AR0803 | *E. faecium* | 12.0 | 48.3 |
| AR0807 | *E. faecium* | 16.0 | 45.8 |
| AR0781 | *E. faecalis* | 13.0 | 98.4 |
| AR0782 | *E. faecalis* | 12.2 | 45.3 |
| AR0801 | *E. casseliflavus* | 11.3 | 45.2 |
| AR0806 | *E. lactis* | 8.5 | 47.1 |

**Table 2.** Plasmids used in this publication.

| **Plasmid** | **Purpose** | **Key Features** | **Selection gene(s)** | **Antibiotic Selection** |
| --- | --- | --- | --- | --- |
| pBRT25-eUnaG | eUnaG expression | *pAMβ1, oriT* | *ant9, cat* | Spectinomycin, chloramphenicol |
| pBRT26-Halo-eUnaG2 | HaloTag9-eUnaG2 expression | *pAMβ1, oriT* | *ant9, cat* | Spectinomycin, chloramphenicol |
| pBRT27-smURFP/eUnaG2 | smURFP/eUnaG2 expression | *pAMβ1, oriT* | *ant9, cat* | Spectinomycin, chloramphenicol |
| pBRT31-smURFP | smURFP expression | *pAMβ1, oriT* | *ant9, cat* | Spectinomycin, chloramphenicol |
| pBRT32-eUnaG2 | eUnaG2 expression | *pAMβ1, oriT* | *ant9, cat* | Spectinomycin, chloramphenicol |
| pBRT38-pac-eUnaG2 | *pac*; eUnaG2 expression | *pAMβ1, oriT* | *pac, cat* | Puromycin, chloramphenicol |
| pBRT48-675-DUAL-gent | 675/680 DUAL smURFP/eUnaG2 genomic insertion | *pWV01ts*, 675/680 homology arms | *aph2, cat, pheS\*\** | Gentamicin, chloramphenicol |
| pBRT63-12650-DUAL-gent | 12650/12655 DUAL smURFP/eUnaG2 genomic insertion | *pWV01ts*, 12650/12655 homology arms | *aph2, cat, pheS\*\** | Gentamicin, chloramphenicol |
| pBRT105-675-eUnaG2 | 675/680 eUnaG2 genomic insertion | *pWV01ts, oriT*, 675/680 homology arms | *aph2, cat, ant9, pheS\*\** | Gentamicin, chloramphenicol, spectinomycin |
| pBRT106-675-2xsmURFP | 675/680 2×smURFP genomic insertion | *pWV01ts, oriT*, 675/680 homology arms | *aph2, cat, ant9, pheS\*\** | Gentamicin, chloramphenicol, spectinomycin |
| pBRT108-12650-eUnaG2 | 12650/12655 eUnaG2 genomic insertion | *pWV01ts, oriT*, 12650/12655 homology arms | *aph2, cat, ant9, pheS\*\** | Gentamicin, chloramphenicol, spectinomycin |
| pBRT109-12650-2xsmURFP | 12650/12655 2×smURFP genomic insertion | *pWV01ts, oriT*, 12650/12655 homology arms | *aph2, cat, ant9, pheS\*\** | Gentamicin, chloramphenicol, spectinomycin |

### Plasmid Transformation

Electrocompetent cells were prepared from overnight cultures diluted 1:1000 into 25 mL BHI containing 200 mM sucrose and glycine (5% [wt/vol] for *E. faecalis*, 1% [wt/vol] for *E. faecium*) and grown overnight. Cultures were pelleted, resuspended in fresh supplemented BHI for 1 h at 37°C, washed three times in ice-cold 500 mM sucrose-10% glycerol, and concentrated to ∼500 µL. For electroporation, 50 µL cells were mixed with 500 ng to 2 µg plasmid DNA, incubated on ice for 30 min, pulsed at 1.8 kV in 1-mm cuvettes, and recovered in BHI plus 500 mM sucrose for 2 h at 37°C or 6 to 8 h at 28°C for temperature-sensitive plasmids before plating on selective BHI agar.

The *E. faecalis* conjugal-delivery strain OG1 derivative, CK111, was employed to transform difficult strains via conjugation(12). CK111 was first transformed via electroporation as above with plasmid DNA that contains the pCF10 conjugal origin. CK111 containing plasmid was grown overnight under selection, washed 2x in fresh media, then CK111 donor was mixed at a ratio of 10:1 with fresh cultures of VREfm recipient. For particularly difficult recipients, the cCF10 peptide pheromone (LVTLVFV) was supplemented at 20 ng/mL prior to plating, which yields >10x more transformants. This mixture was incubated overnight on filter paper on BHI agar without selection. Filter paper matings were then removed and resuspend in BHI prior to plating on agar containing 20 µg/mL vancomycin and appropriate selection antibiotics.

### Minimum inhibitory concentration (MIC) assay

MICs were determined by broth microdilution. Overnight cultures were adjusted to ∼5 × 10^5 CFU/mL in fresh BHI and added to 2-fold antibiotic dilutions (0.8 to 400 µg/mL) in flat-bottom 96-well plates. Negative controls contained BHI alone; positive controls contained bacteria without antibiotic. Plates were incubated aerobically at 37°C for 18 to 24 h. MICs were defined as the lowest concentration preventing visible growth, based on OD600 and visual confirmation. Assays were performed in biological triplicate.

### Genomic Integration

To generate stable genomic insertions, smURFP/eUnaG2 reporter plasmids containing homology to the 675/680 chromosomal region or the 12650/12655 chromosomal region were transformed into the VREfm strain as described above, and single colonies were picked and maintained at 28°C under dual selection (gentamicin/chloramphenicol or gentamicin/spectinomycin) for one passage. Next cultures were subcultured twice at 1:1000 in BHI containing only 300 µg/mL gentamicin at 37°C to encourage recombination and plasmid loss. Finally, cultures were serially diluted and plated onto BHI supplemented with 300 µg/mL gentamicin and 5-10mM PCPA. Successful integration was confirmed by screening single colonies for genomic insertion by PCR. (675-detect-F, 5’-catgacaagtaaggacagctatatcgtccc ; 680-detect-R, 5’-cgtggcaaacgagtcactagcaag ; 12650-detect-F, 5’-cgtttatagattggctgtggctttg ; 12655-detect-R, 5’-ctcactgaaagaacaactcgcatc).

### Anaerobic Reporter Fluorescence

VREfm transformants expressing HaloTag9, eUnaG, eUnaG2, or smURFP from the *S. agalactiae cfb* promoter were grown in BHI at 37°C. HaloTag labeling used 1 µM HaloTag TMR ligand for 1 h at 25°C followed by 2 to 3 PBS washes. eUnaG and eUnaG2 cultures were supplemented with 12.5 to 25 nM bilirubin, and smURFP cultures with 2.5 to 5 nM biliverdin, during overnight growth. Because BHI has background fluorescence, cells were pelleted and resuspended in PBS before measurement. Unlike HaloTag9, eUnaG2 and smURFP require ligand supplementation only *in vitro* and do not require wash steps(18).

### Mouse colonization model

Six- to eight-week-old female C57BL/6NTac mice (Taconic) received vancomycin hydrochloride oral solution (0.5 mg/mL) in drinking water for 2 days before inoculation with VREfm. Mice were gavaged with 1×10^8^ CFU of indicated VREfm for reporter plasmid-containing strains or 1×10⁴ CFU VREfm for other strains and monitored by plating fresh stool. Stool was resuspended at 10% (wt/vol) in sterile PBS, and where appropriate was serially diluted before plating on BHI agar containing vancomycin (30 µg/mL) and meropenem (20 µg/mL).

### Ethics statement

All animal experiments were approved by the Institutional Animal Care and Use Committee of St. Jude Children’s Research Hospital under protocol 3142 and were performed in accordance with institutional guidelines and applicable regulations.

### Flow cytometry

Fresh stool was collected and was resuspended at 10% (wt/vol) in PBS, filtered through 40-µm strainers, diluted 10-fold in 1XPBS containing 2.5 nM biliverdin and 12.5 nM bilirubin, incubated on ice for one hour, spun and resuspended in fresh 1xPBS, then analyzed on an ApogeeFlow MicroPLUS. SALS and LALS were collected with a 405-nm laser. eUnaG2 was detected at 488/530-40 and smURFP at 638/676-36. Data were analyzed in FlowJo v10.10. For Figure 3A and Supplemental Figure 4B, events were first gated broadly on the bacterial population using SALS and LALS. For Supplemental Figure 5, a narrower VRE gate was defined using VRE-only controls before singlet and fluorescence gating.

### Fluorescent Imaging

Tissues were embedded in tissue freezing medium, snap-frozen in liquid-nitrogen-cooled isopentane, sectioned at 10 µm, and dried for 1 h at room temperature. Sections were rehydrated in PBS, blocked in PBS with 1% (wt/vol) BSA and 0.05% Tween-20, stained with wheat germ agglutinin-Alexa Fluor 555 (1 µg/mL) for 1 h, washed, and mounted with Vectashield containing DAPI. Images were acquired on a Nikon Ti2 with a 60× Plan Apo oil objective, Lumencor Spectra/AuraII illumination, and Hamamatsu Orca Fusion camera. Z-stacks were collected at 0.2-µm steps and processed in NIS-Elements v5.42.04.

### *In vitro* competition assay

For *in vitro* competition assays, VREfm strains carrying genomic insertions of either eUnaG2 or 2×smURFP were grown overnight, adjusted to equal input amounts, and mixed at 10^4 CFU per strain in BHI supplemented with biliverdin (5 nM) and bilirubin (25 nM) in a total volume of 200 µL in 96-well plates. Plates were incubated anaerobically for 16 h, after which cultures were pelleted, resuspended in 1× PBS, and fluorescence and OD₆₀₀ were measured. Fluorescence values were first normalized to OD₆₀₀ and then scaled relative to matched single-color controls to estimate the relative contribution of each strain to the mixed culture. A relative competition index was calculated as the log2-transformed output ratio, log2(smURFP/eUnaG2), where positive values indicate relative enrichment of the 2×smURFP-labeled strain and negative values indicate relative enrichment of the eUnaG2-labeled strain.

## Acknowledgements

We would like to thank Daria van Tyne (University of Pittsburgh) and Jason Rosch (St. Jude Children’s Research Hospital) for protocols, strains(32), and helpful advice in working with difficult VREfm. CK111, the conjugal donor *E. faecalis* strain(12), as well as several plasmids were a kind gift of Danielle Garsin. Plasmid pBSU101-EGFP, which was the basis for anaerobic reporter plasmid constructs, was a kind gift from Kevin Wood(21, 33). Plasmid pMAL-c5x_UnaG was a gift from Julie Biteen (Addgene#163125). Plasmid pJYDN3p was a gift from Gideon Schreiber (Addgene#162457). Plasmid pBAD-smURFP-RBS-HO-1 was a gift from Erik Rodriguez and Roger Tsien (Addgene# 80341). Plasmid pET51b(+)HaloTag9 was a gift from Kai Johnsson (Addgene#169324). Plasmid pGPA1, which was the basis of the temperature-sensitive genomic insertion plasmids, was a gift from Willem van Schaik (Addgene#115476). This work was supported by The American Lebanese Syrian Associated Charities (ALSAC) and by the National Institute of Health (NIH) grant R21AI180489-01.

## Author contributions

B.R.T. conceived the study, designed and performed experiments, analyzed data, and wrote the original draft. J.H., M.T., C.G., C.B., M.D., T.F., M.B., and C.M.K. contributed to investigation, methodology, reagents, or data analysis and reviewed and edited the manuscript. E.B.M. conceived and supervised the study, acquired funding, and reviewed and edited the manuscript. All authors approved the final manuscript.

## Data availability

All data supporting the findings of this study are included in the article and its supplemental material. Plasmids described in this study will be deposited with Addgene prior to publication. Additional materials are available from the corresponding author upon reasonable request.

