## Supplemental Data for "Anaerobic Fluorescent Reporters Enable *In Vivo* Tracking of VRE Dynamics in *Enterococcus faecium*"

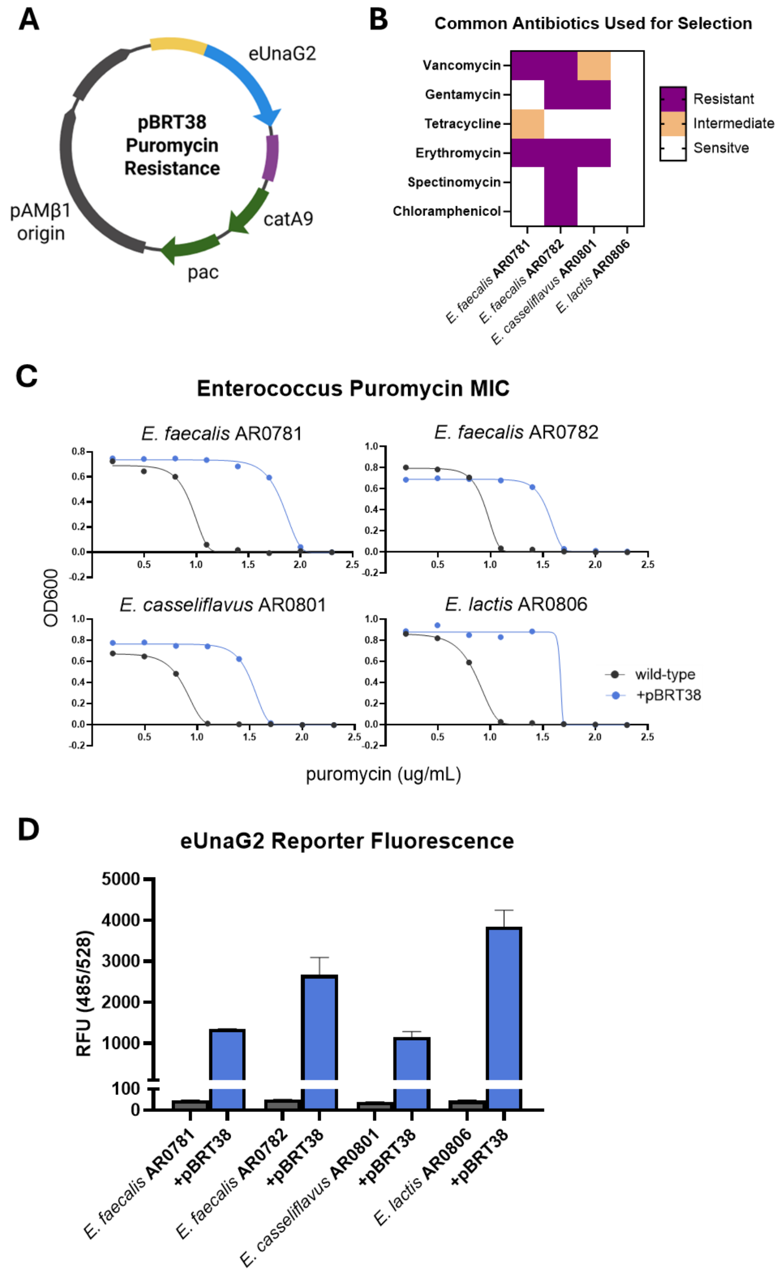


**Supplemental Figure 1. Puromycin-Resistance Marker *pac* is Broadly Compatible with Enterococci.** (**A**) Plasmid schematic of pBRT38 plasmid containing eUnaG2 reporter, a chloramphenicol resistance gene, a codon-optimized *pac* gene under control of a duplicate chloramphenicol promoter. (**B**) Heatmap of resistances to many common antibiotics for selection of 4 non-faecium Enterococcus strains from CDC AR Isolate Bank,. (**C**) MIC assays for puromycin of 4 Enterococcus strains, wild-type are in blue, and transformants expressing *pac* from pBRT38 are in black. (**D**) Relative fluorescence of strains transformed with puromycin-resistant eUnaG2 reporter plasmids after overnight growth in BHI supplemented with 25nM bilirubin.

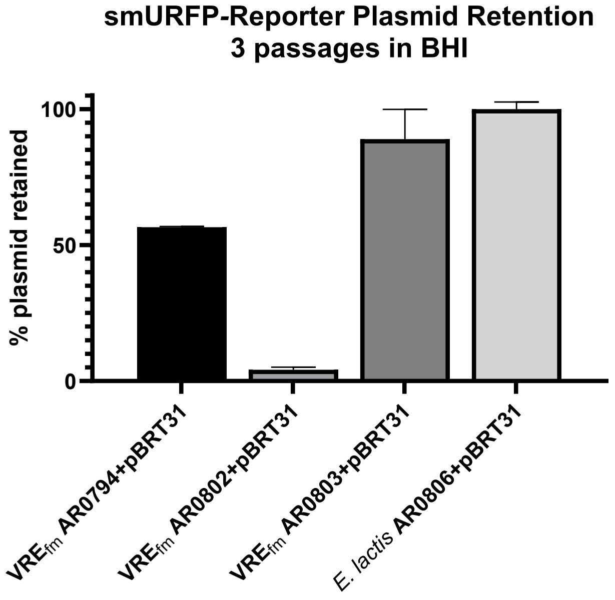


**Supplemental Figure 2. Plasmid Instability in VREfm.** Four *E. faeciu*m strains from the CDC AR Isolate Bank were transformed with plasmid pBRT31 (smURFP expressing), maintained under strict selection (500 µg/mL spectinomycin, 30 µg/mL chloramphenicol) prior to passaging (1:1000) single colony isolates 3 times in BHI without selection. Cultures were then serially diluted and plated on BHI agar alone or BHI agar supplemented with 500 µg/mL spectinomycin and 30 µg/mL chloramphenicol. Colonies were enumerated and the percent plasmid retention calculated. Assays were performed in triplicate, and the percentage of colonies that retained resistance plotted as a mean with S.D.

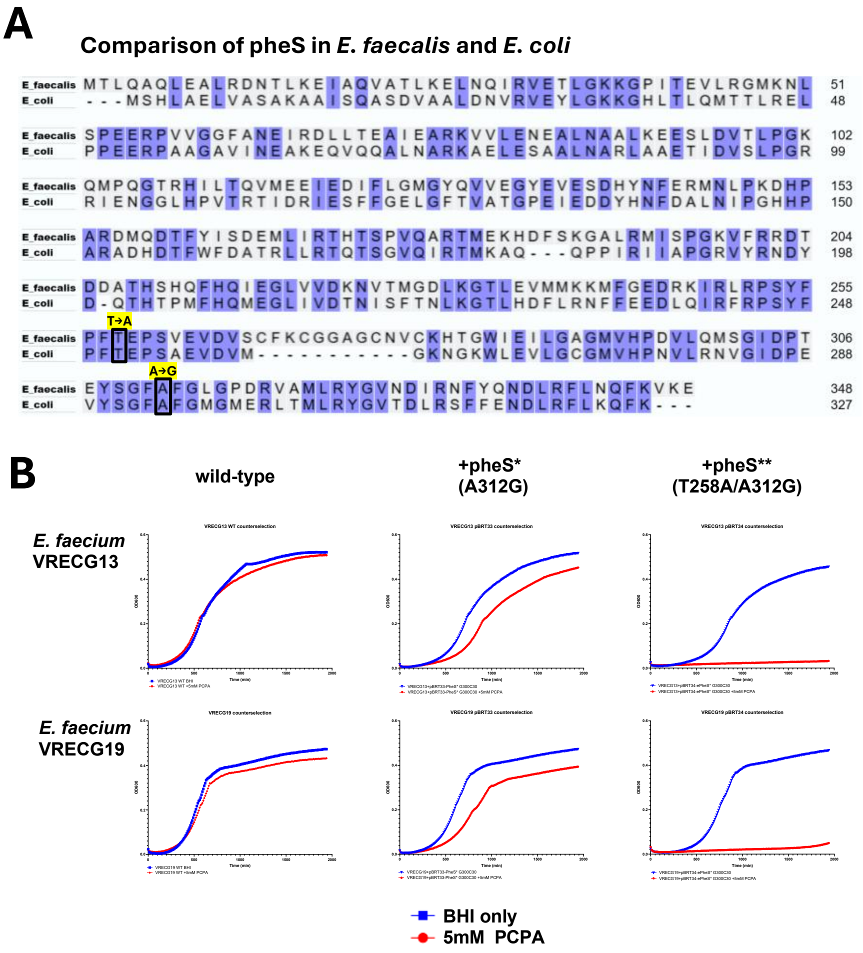


**Supplemental Figure 3. Improved PheS** Counterselection in *E. faecium*** (**A**) Partial amino acid sequence comparison of *E. coli* and *E. faecalis pheS*, highlighting mutations to generate *pheS**(A312G) and *pheS***(T258A/A312G) counterselection genes. (**B**) Growth curves of three *E. faecium* strains of wild-type and transformants containing plasmid pBRT33 (*pheS**) or pBRT34 (*pheS***). Transformants were grown in media containing BHI only or 5 mM p-chlorophenylalanine (PCPA). Sequence alignment was generated using the UniProt Align tool[34].

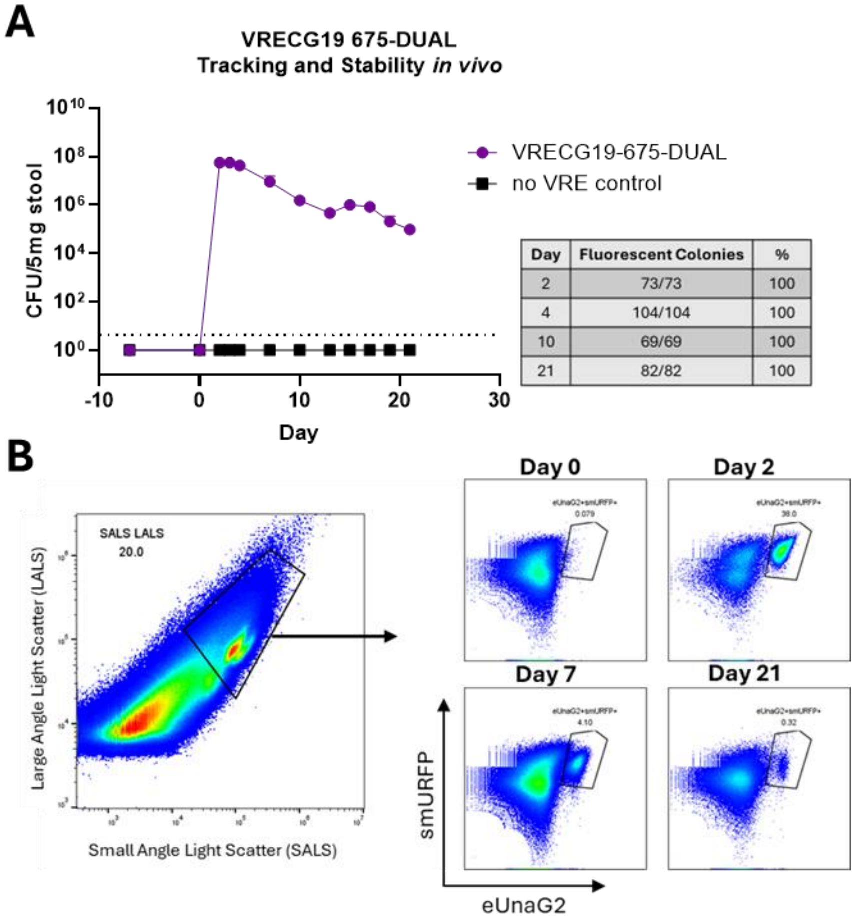


**Supplemental Figure 4. Tracking VRECG19-675-DUAL *In Vivo*** (**A**) Female C57BL/6 mice (6–8 weeks; n=5/group, Taconic Biosciences) received vancomycin (0.5 mg/mL in drinking water) for 7 days before inoculation with $1\times10^8$ CFU E. faecium VRECG19-675-DUAL. Stool collected at indicated days post-inoculation (dpi) was resuspended in PBS (10% w/v), serially diluted, and plated on BHI agar supplemented with vancomycin (30 µg/mL), meropenem (20 µg/mL), and biliverdin (5 nM). Colony fluorescence was confirmed by fluorescence imaging of whole plates (BioRad ChemiDoc MP; 647 nm excitation, 700/50 filter) and fluorescent CFU enumerated at 2, 4, 10, and 21 dpi. (**B**) Stool from VRECG19-675 DUAL inoculated mice was resuspended in PBS (10% w/v) containing biliverdin (2.5 nM) and bilirubin (12.5 nM), incubated anaerobically on ice for 30 min, centrifuged through a 40 µm mesh filter, and resuspended. Flow cytometry was performed on an ApogeeFlow Micro-PLUS. Events were gated broadly on the total bacterial population using SALS and LALS parameters, before fluorescence gating for smURFP and eUnaG2.

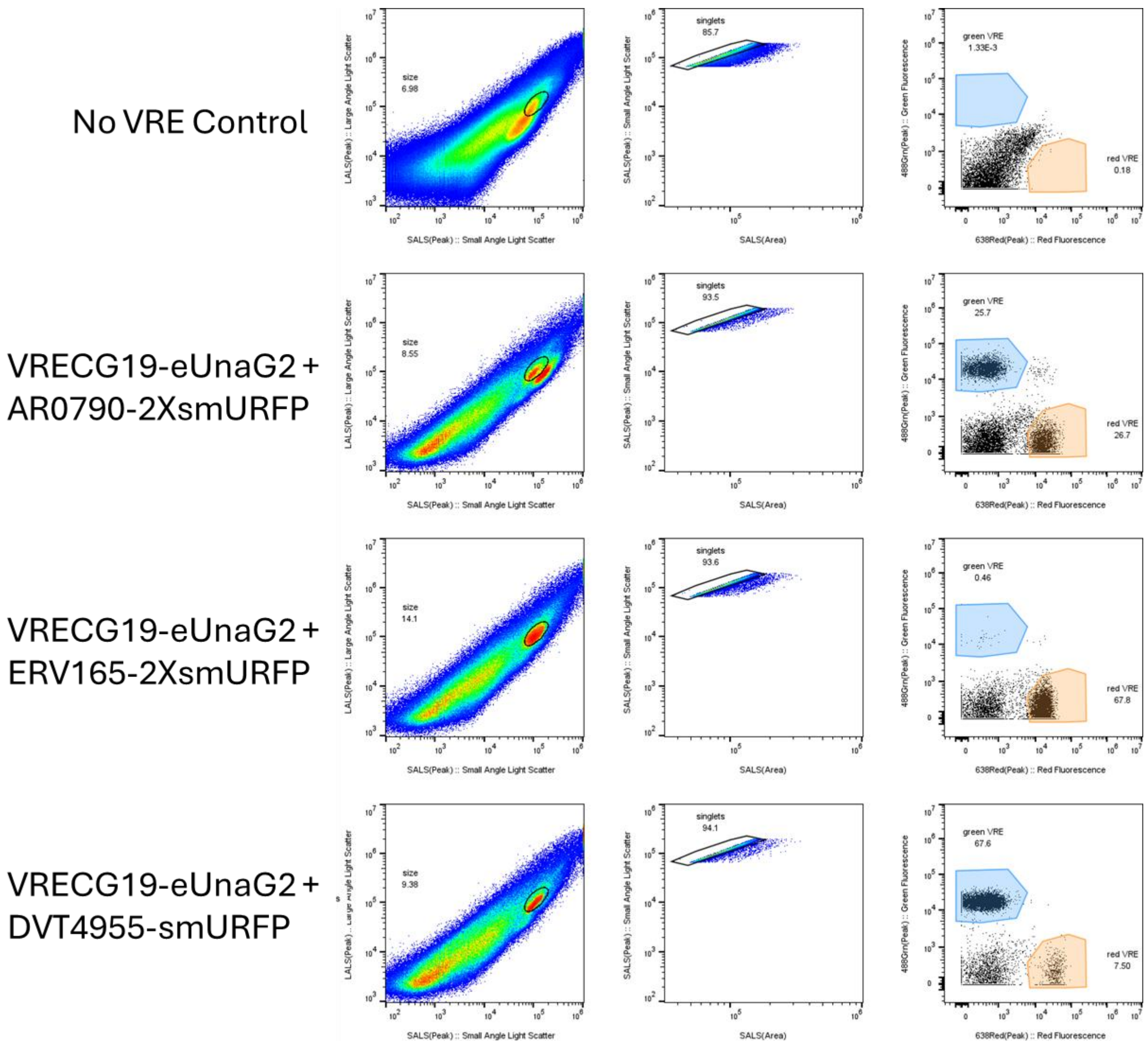


**Supplemental Figure 5. Flow Cytometry Gating for *in vivo* Competition.** Female C57BL/6 mice (6–8 weeks) received vancomycin (0.5 mg/mL) for 48 h before gavage with equal mixtures of eUnaG2- and 2×smURFP-expressing VREfm ($10^4$ CFU each). Stool samples were analyzed by flow cytometry to quantify eUnaG2$^+$ and smURFP$^+$ VREfm. Fresh stool (1–21 dpi) was resuspended in PBS (10% w/v), filtered through 40 µm strainers, diluted 10× in PBS, and 50 µL analyzed on an ApogeeFlow Micro-PLUS. SALS and LALS (forward/side scatter equivalents) were detected with a 405 nm laser. eUnaG2 fluorescence was detected with a 488 nm laser and 530/40 bandpass filter, and smURFP with a 638 nm laser and 676/36 bandpass filter. Events were first gated using a narrow VRE gate defined from VRE-only controls in LALS vs. SALS space, followed by singlet gating (SALS peak vs. SALS area) and fluorescence gating of eUnaG2$^+$ and smURFP$^+$ populations using single-color controls.
